# Epigenetic age estimation of wild mice using faecal samples

**DOI:** 10.1101/2023.10.25.563913

**Authors:** Eveliina Hanski, Susan Joseph, Aura Raulo, Klara M Wanelik, Áine O’Toole, Sarah C L Knowles, Tom J Little

## Abstract

Age is a key parameter in population ecology, with a myriad of biological processes changing with age as organisms develop in early life then later senesce. As age is often hard to accurately measure with non-lethal methods, epigenetic methods of age estimation (epigenetic clocks) have become a popular tool in animal ecology, and are often developed or calibrated using captive animals of known age. However, studies typically rely on invasive blood or tissue samples, which limits their application in more sensitive or elusive species. Moreover, few studies have directly assessed how methylation patterns and epigenetic age estimates compare across environmental contexts (e.g., captive or lab-based vs. wild animals). Here, we built a targeted epigenetic clock from laboratory house mice (strain C57BL/6, *Mus musculus)* using DNA from non-invasive faecal samples, and then used it to estimate age in a population of wild mice (*Mus musculus domesticus*) of unknown age. This lab mouse-derived epigenetic clock accurately predicted adult wild mice to be older than juveniles, and showed that wild mice typically increased in epigenetic age over time, but with wide variation in epigenetic ageing rate among individuals. Our results also suggested that, for a given body mass, wild mice had higher methylation across targeted CpG sites than lab mice (and consistently higher epigenetic age estimates as a result), even among the smallest, juvenile mice. This suggests wild and lab mice may display different CpG methylation levels from very early in life, and indicates caution is needed when developing epigenetic clocks on lab animals and applying them in the wild.

## Introduction

Age is a key characteristic for any organism, with numerous biological processes from immune maturation to reproduction being age-related (Georgountzou & Papadopoulos, 2017). Yet, measuring age in wild individuals can be challenging as date of birth is often unknown. Classic methods for assessing age in wild animals are often inaccurate (e.g., body size) or destructive (e.g., eye lens weight), raising ethical concerns and preventing longitudinal studies. An alternative approach for measuring the age of wild individuals relies on the measurement of epigenetic marks in particular genomic regions. Specifically, at some CpG sites (cytosines followed by a guanine; Moore et al., 2013) the proportion of methylated cytosines appears to change linearly with age. DNA methylation can influence biological ageing through various molecular mechanisms, including repression of chromatin state and promotor silencing among others (Moore et al., 2013). Together these CpG sites with age-related methylation patterns can be used to derive an ‘epigenetic clock’. An epigenetic clock, trained using samples from individuals of known age, can then be used to predict age in individuals of unknown age. Such epigenetic clocks can provide a more accurate estimate of chronological age among wild animals than visible characteristics (Mayne et al., 2022; Larison et al., 2021). Epigenetic clocks have now been developed for a wide range of animal species including baboons, chimpanzees, humpback whales, wolves, green turtles, and zebras (Jarman et al., 2015; Anderson et al., 2021; Pinho et al., 2022; De Paoli-Iseppi et al., 2017; Polanowski et al., 2014; Thompson et al., 2017; Wright et al., 2018; Mayne et al., 2022; Larison et al., 2021; Bors et al., 2021; Sullivan et al., 2022; Tangili et al., 2023; Ito et al., 2018; Fairfield et al., 2021; Wilkinson et al., 2021), as well as plants (Gardner et al., 2023).

Alongside measuring chronological age, epigenetic clocks also appear to capture signals of biological age, typically considered to reflect the accumulated damage and functional decline in cells, tissues, and organs (Yousefzadeh et al., 2021). Accelerated epigenetic ageing has been linked to various communicable and non-communicable diseases in both humans and laboratory mice (Joyce et al., 2021; Morales Berstein et al., 2022; Ambatipudi et al., 2017; Harvanek et al., 2021; Cao et al., 2022; Peng et al., 2019). Insights have also come from the wild: high social rank is associated with accelerated epigenetic ageing in wild baboons (Anderson et al., 2021), and hibernation slows down ageing in marmots and bats (Pinho et al., 2022; Sullivan et al., 2022). Thus, the use of epigenetic clocks may provide a means of estimating chronological age among wild animals while simultaneously providing insight into biological ageing in natural settings.

Previous studies using epigenetic clocks have focused on humans (Joyce et al., 2021; Morales Berstein et al., 2022; Ambatipudi et al., 2017; Harvanek et al., 2021; Cao et al., 2022; Peng et al., 2019), laboratory (Han et al., 2018; Kerepesi et al., 2022) or wild animals (Prado et al., 2021; Polanowski et al., 2014; Anderson et al., 2021; De Paoli-Iseppi et al., 2018; Lemaître et al., 2022). A number of studies have also included both wild and captive (e.g., from a zoo) individuals (Mayne et al., 2022; Robeck et al., 2021; Ito et al., 2018; Fairfield et al., 2021; Wilkinson et al., 2021). However, a comparison of wild and laboratory individuals has not been previously conducted. Comparing epigenetic age and ageing between lab and wild individuals of the same species could help us understand drivers of biological ageing and its variability in individuals from contrasting genetic and environmental backgrounds.

Here, we build an epigenetic clock using samples from laboratory mice (*Mus musculus*) and use this lab-based clock to predict age in house mice (*Mus musculus domesticus*) from a wild population. Our aims were threefold: (1) to see whether we could develop an epigenetic clock from lab-based animals capable of accurately capturing differences in chronological age within a wild population, (2) to compare estimates of epigenetic age in lab compared to wild mice of a given size, to gain insight into biological ageing in lab vs wild settings, and (3) to assess the extent of variability in biological (epigenetic) ageing rate among wild mice. We used faecal samples as a source of DNA, to develop a non-invasive method that allows longitudinal sampling without ethical or logistical limitations on sampling frequency, and allows applications of epigenetic age estimates in contexts where animal capture or handling are not possible. To our knowledge, this is the first epigenetic clock built with faecal samples. Our results show the potential of such an approach, but also indicate substantial differences in DNA methylation levels of an inbred laboratory population and an outbred wild population, even from very early in life.

## Materials and Methods

### Sample collection

A total of 137 faecal samples were collected from 65 individual *Mus musculus* C57BL/6 laboratory mice (30 females, 35 males) from two animal facilities. The samples were collected in May– November 2021 at the Biomedical Services Building, Oxford, UK (Animal facility A), and King’s College, London, UK (Animal facility B). The chronological age of the mice varied from 7 to 339 days, covering approximately the first third of expected C57BL/6 lifespan (30–32 months; Schultz et al., 2020). The mice were kept in standard housing and were not subject to any interventions before or during sampling. During sample collection, body mass was recorded for mice from Animal facility B but not for mice from Animal facility A. However, body mass is tightly correlated with age among juvenile house mice (Spangenberg et al., 2014; Jax, 2022a; Jax, 2022b) allowing accurate estimation of mass from age. As such, we estimated body mass for 25 mice under seven weeks of age from Animal Facility A (for older mice from this facility (*n*=8) body mass was not estimated and consequently samples from those mice were not included in analyses including body mass). Body mass estimation was done based on Spangenberg et al. (2014) for 7– 20-day old pups and The Jackson Laboratory C57BL/6 body mass references for 3–7-week-old pups (Jax, 2022a; Jax, 2022b). For the latter age group, estimation was done separately for females and males using the Jackson Laboratory sex-specific data (Jax, 2022a; Jax, 2022b). Mice were weighed by placing them on a scale. To collect faecal samples, mice were briefly placed on a sterile surface until defecation. Faecal pellets were collected in a sterile manner, immediately preserved in DNA/RNA Shield, and stored frozen at −80°C until further processing (up to ≤12 months).

Wild house mouse (*Mus musculus domesticus*) sampling was conducted in April–May 2019, July 2019, September–October 2019, August–September 2020, and April–May 2021 on Skokholm Island, Wales, UK. Mice were trapped overnight using small galvanised metal Sherman live traps baited with peanuts and non-absorbent cotton wool for bedding, and with a spray of sesame oil outside the trap as a lure. Across each of two broad sampling areas (one near the coast and one in the island interior, named “Quarry” and “Observatory” respectively), on each trapping night 150 traps were set at dusk and checked at dawn. To prevent cross-contamination, any traps showing signs of mouse presence were washed and sterilised before being reset using bleach solution (including at least a 60 min soak in 20% bleach solution) to destroy bacterial cells and DNA. All newly captured mice were permanently identified by subcutaneous injection of a passive integrated transponder (PIT) tag. Upon each capture, each mouse was either tagged or identified (if a recapture), aged, sexed, and measured before being released within 3m of its trapping point. Sex was determined using anogenital distance and reproductive state. Reproductive state was recorded as either non-perforate, perforate, suspected pregnant, or lactating for *females*, and testes abdominal, small or large for *males*. At each capture, body mass was recorded. Mice were placed in a small cotton bag for weighing to the nearest 0.1g, and in a transparent plastic bag for measurement of body length (measured as snout-vent length, SVL, from the tip of the nose to the base of the tail, with mice gently straightened before measuring). Reproductive state was classified as active or inactive, with females being reproductively active when either pregnant, lactating, or perforate, and males being reproductively active when testes were visibly descended. Age was roughly classified to one of three age groups using body mass and reproductive state. Reproductively inactive mice weighing ≤15.0g were classified as *juveniles*, mice >20.0g of body mass were classified as *adults* regardless of reproductive state, and mice falling between these two categories (reproductively active mice ≤15.0g of body mass, as well as all mice weighing 15.1– 20.0g) were classed as *sub-adults*.

Faecal samples were collected from traps in a sterile manner (shortly after mice were collected from traps the following day), preserved in DNA/RNA Shield and stored in a −20°C freezer until the end of fieldwork (maximum 6 weeks after sample collection). At this point they were returned to the laboratory frozen and stored at −80°C until DNA extraction (up to ≤17 months). A total of 215 samples were selected from all collected samples (>900) for further processing, by first selecting a longitudinal dataset (mice sampled ≥2 times over time; total of 54 individuals) with as much variation as possible in morphometric variables (age, body mass, sex, and reproductive status), as well as environmental variables (sampling season and sampling area) and then supplementing this with additional (equally variable) cross-sectional samples (one sample per animal) to increase number of individuals for cross-sectional analyses up to 130. Variation in variables was achieved by randomly selecting approximately equal numbers of samples across categories e.g., across juvenile, sub-adult and adult mice.

### DNA extraction, bisulfite conversion and PCR amplification

DNA was extracted from faecal samples using the ZymoBIOMICS DNA MiniPrep Kit according to the manufacturer’s protocol (Zymo Research, Irvine, California, USA). DNA was then bisulfite-converted using the Zymo EZ DNA Methylation-Gold Kit to convert unmethylated cytosines to uracil and then thymine (Zymo Research, Irvine, California, USA). PCR amplification was conducted for five genes previously reported to correlate with chronological age in *Mus musculus*; *Prima1, Hsf4, Kcns1, Gm9312*, and *Gm7325* (Han et al., 2018). Amplification was conducted using the PyroMark PCR Kit according to manufacturer’s instructions and primers for the five genes (QIAGEN, Hilden, Germany; Table S2; Han et al., 2018). PCR conditions were as follows: 15 min initial denaturation at 95℃, 50 cycles of 30 sec denaturation at 58℃, 30 sec primer annealing at 58℃ and 30 sec extension at 72℃, followed by a 10 minute final extension at 72℃. Amplification success was confirmed using gel electrophoresis. PCR was repeated with the same conditions for any reaction that did not produce a band on the gel. The five amplicons (PCR products) were pooled for each sample into 5-gene libraries by combining all PCR products per sample. DNA was quantified with Qubit Fluorometer High Sensitivity dsDNA kit and normalised to 6.25 ng/*μ*l (Thermo Fisher Scientific, Waltham, Massachusetts, USA).

### Sequencing, basecalling, and demultiplexing

The Oxford Nanopore Technology (ONT) platform was used for library sequencing, and all ONT procedures were conducted according to manufacturer’s instructions and ONT protocol NBA_9102_v109_revl_09Jul2020 (Oxford Nanopore Technologies, Oxford, UK). We first used the ONT Ligation Sequencing Kit (SQK-LSK109) to repair and dA-tail the DNA ends, followed by ligation of sequencing adaptors to the prepared ends. We then barcoded libraries using the ONT Native Barcoding Expansion kit (EXP-NBD104 or EXP-NBD196). Approximately 15 ng of the prepared library was loaded onto a prepared ONT MinION Mk1B R9.4.1 flow cell and sequenced using the ONT MinKNOW software v21.10.4. Libraries were sequenced across a total of six runs resulting in a mean of 49,969 reads per sample. A negative control (where DNAse free H_2_O was used instead of pooled amplicons at the start of Nanopore pipeline) was included in three sequencing runs, and these generated a mean of 200 (range 17–519) reads. One flow cell was used twice, and washed between runs with the ONT Flow Cell Wash kit (EXP-WSH003). Different barcodes were used for negative controls across the two sequencing runs where the same flow cell was used to enable testing for carry-over of reads (only 17 potential carry-over reads were detected). Raw sequencing data was basecalled and demultiplexed using High Accuracy basecalling on the ONT Guppy software v5.0.11. The basecalled FASTQ files were then run through the Apollo pipeline v0.1 (https://github.com/WildANimalClocks/apollo, DOI: 10.5281/zenodo.8426692) to acquire methylation rates for each CpG site within the five genes (73 CpG sites in total, 4–27 CpG sites per gene). Apollo requires a minimum read count of 50 for reporting on a particular site. Using the alignment with the reference genes, target sites (cytosines within CpGs) were identified in each read and determined as either methylated (cytosine) or unmethylated (uracil). The process was continued for each read, resulting in a proportion of methylated cytosines at each CpG site.

### Analyses

The data was analysed and visualised in R v4.1.2 (R Core Team, 2023). The epigenetic clock was built using the cross-sectional data of 50 samples from C57BL/6 mice housed in two facilities (Table S1). Using CpG site specific methylation rates, we used the package glmnet v4.1–3 (Friedman et al., 2010) to perform a cross-validated elastic net regularisation using the cv.glmnet function using a LASSO model (mixing parameter alpha=1) and a leave-one-out cross validation (nfolds=nrow). From inspecting relationships between methylation levels in CpG sites and body mass (Supplementary Figure 1–4), we expected only a subset of CpG sites to be relevant for predicting age and thus alpha parameter was set to 1 (LASSO penalty). We further investigated the effect of different alpha values on model fit by changing alpha between 0 (ridge penalty) and 1 (LASSO penalty) at increments of 0.05 and selecting a value of alpha that maximises model fit (defined by lowest mean squared error). This analysis was ran 10 times. The analyses did not suggest a strong lead candidate for alpha value; however, our initial alpha value of 1 was amongst the strongest candidates and as such we proceeded with alpha=1. We then fitted a final glmnet model using optimal lambda value determined by the cross-validation (lambda=0.913). Epigenetic age was then predicted based on the glmnet model using the predict() function. This clock was validated on an additional 15 C57BL/6 mice from Facility B (Table S1). Using one sample per animal, we first measured correlation between epigenetic age and chronological age, and assessed clock performance for estimating chronological age of lab mice using mean absolute error (MAE; Tangili et al., 2023). We then used a linear mixed effects model using lmer function from lme4 R package (Bates et al., 2015) to measure the influence of sex and sequencing batch on epigenetic age predictions. For this model we used two samples from each 15 mice (total 30 samples) and included animal ID as a random factor.

The above-described epigenetic clock (clock 1) was developed to test and showcase how accurate a faeces-based targeted epigenetic clock can be. However, as our aim was to develop a lab-based epigenetic clock that could be used to predict age in wild mice, we then built a second epigenetic clock (Clock 2) as described above but using only those CpG sites that (1) showed parallel trends in their methylation rates against body mass in lab and wild mice (i.e., ones that increased/decreased in methylation with body mass in both systems based on linear regression slope estimates quantifying the change in DNA methylation levels per unit change in body mass for specific CpG sites; Fig. S1–4), to account for lab vs wild changes in methylation that may be driven by genetic and environmental differences, and (2) had methylation rates for the majority of wild mouse samples (we failed to acquire sufficient read counts for methylation rate measurement for all 11 CpG sites from the gene *Gm7325* in 9% of all wild mouse samples). Following these principles, we included a total of 53 CpG sites in Clock 2. Lambda value used in this clock (determined with cross-validation, as for Clock 1) was 16.950. Clock 2 was similarly validated with the independent lab dataset and then used to estimate epigenetic age for all wild house mouse samples for which methylation rates at the CpG sites included in the clock were successfully measured (*n*=201; 93% of all wild mouse samples; Table S1) using linear modelling. Intercepts and coefficients for both epigenetic clocks developed are presented in Table S5.

To examine method repeatability, 11 out of the 201 wild mouse DNA samples were processed twice through BS treatment, PCR and sequencing. First and repeat DNA aliquots were processed in two distinct batches. Repeatability was estimated for (1) methylation levels of CpG sites included in the clock 2 (*n*=11; Table S4) and (2) epigenetic age estimates, using R package rptR (Stoffel et al., 2017) with 1,000 parametric bootstraps. Sequencing batch and sample ID were used as predictors (random effects). After assessing repeatability, duplicates were removed from the data by randomly selecting one observation per sample ID.

To test for the effect of covariates on predicted epigenetic age in the lab mouse validation dataset, we fit a linear model with epigenetic age as the dependent variable and chronological age, sex, cage, and sequencing run ID as predictor variables. ANOVA was used to test whether predicted epigenetic age varies significantly by wild mouse age categories (juvenile/sub-adult/adult) and post-hoc Wilcoxon rank sum tests were used to test whether the predicted epigenetic age of wild mice varied significantly between specific age category pairs (juvenile vs sub-adult, juvenile vs adult, sub-adult vs adult). The ability of the clock to detect an increase in age among wild mice sampled on two consecutive occasions was tested using a one-tailed binomial test. The null hypothesis for this test was that the probability of mice increasing in epigenetic age between consecutive timepoints (p) = 0.5 (i.e., mice are just as likely to increase or decrease in epigenetic age over time), while the alternative hypothesis was that p>0.5. We also used linear models to test (1) whether among wild mice, time between sampling points predicted absolute change in epigenetic age as well as (2) whether sex, trapping area, season (*spring*: April/May; *summer*: July/August; *autumn:* September/October), reproductive activity and body mass as measured at first capture predict rate of epigenetic ageing. Mice with less than 27 days between time-points were excluded from this longitudinal analysis as the mean absolute error (MAE) of the clock in the validation dataset was 26 days (Fig. 1B).

**Figure 1.**
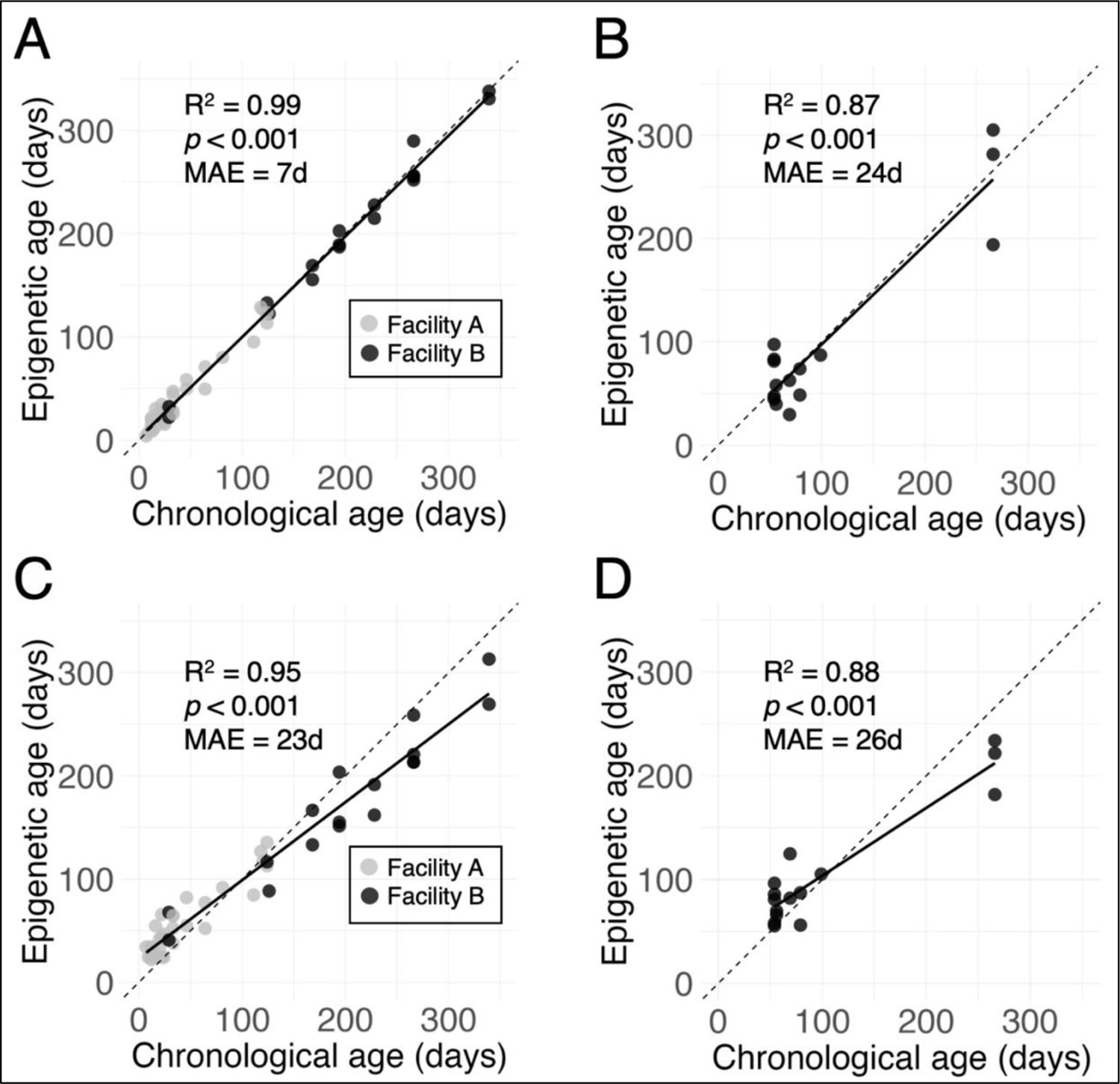
The relationship between DNA methylation-based (epigenetic) age and chronological age in C57BL/6 mice. (**A**, **C**) used to train the epigenetic clock model or (**B**, **D**) an independent set of mice (a validation set) not used in building the epigenetic clock model. Two epigenetic clocks were built: one with 22 CpG sites from five genes (**A**, **B**), and one with 11 CpG sites from two genes (**C**, **D**). Circles represent individual mice, circle colour in A indicates animal facility (*grey*= facility A, *black* = facility B). Solid lines are linear regression lines, while dashed lines are a reference line of *y=x* (the hypothetical relationship if chronological age and epigenetic age estimates were exactly equivalent).

To explore whether lab and wild mice might differ in methylation levels across CpG sites from targeted genes that showed parallel trends in lab and wild (i.e., genes that decreased/increased in methylation with body mass in both systems: *Hsp4, Gm9312, Kcns1, Gm7325;* gene *Prima1* was excluded since it increased in methylation in wild mice but decreased in lab mice), we used body mass as a proxy of age. While the reliability of body mass as an indicator of age declines after initial growth during first few weeks of life, it continues to increase with chronological age in both C57BL/6 lab and wild mice beyond this time and thus can be used as a rough estimate of age in adults as well (Fig. S5; Jax, 2022a; Jax, 2022b; Gerber et al., 2021; Gray et al., 2015). We used a Bayesian regression model brm using R package brms (Bürkner, 2017) with methylation level as response variable and source (lab/wild) and body mass as predictors. The model also included an animal ID random effect since the data used contained ≥1 samples per individual, as well as a nested random effect Gene/Position to account for measurement of methylation at multiple positions within targeted genes. Further, we used a generalized additive model to test whether source (lab/wild) predicted epigenetic age. The following models were compared to explore potential non-linearities in the relationship between epigenetic age and body mass, while seeing whether there are differences in this relationship by source: (1) gam(Epigenetic age ∼ Body mass), (2) gam(Epigenetic age ∼ s(Body mass), (3) gam(Epigenetic age ∼ Body mass + Source), (4) gam(Epigenetic age ∼ s(Body mass) + Source, (5) gam(Epigenetic age ∼ Body mass * Source), and (6) gam(Epigenetic age + s(Body mass, by = Source). Model 5 (gam(Epigenetic age ∼ Body mass * Source)) had the best model fit (assessed from GCV, AIC and adjusted R-squared values) and was then used to test whether source predicts epigenetic age. Female wild mice with signs of ongoing or recent pregnancy (those recorded as suspected pregnant, n=9) were excluded from these models including body mass, as body mass will be a less accurate age proxy in these individuals, who are significantly heavier than other females (one sample per Animal ID, *n*=48; linear model, *F*_1,54_= 29.28, *p*<0.001).

Lastly, to explore whether the *rate* of epigenetic ageing differed between lab and wild mice we used two methods. First, we tested whether the mean rate of epigenetic aging differed between lab and wild mice. To do this we constructed a linear model using data from repeat-sampled mice, where rate of epigenetic ageing was the response, and source (lab/wild) was the predictor, while including body mass and sex as covariates. All repeat-sampled lab mice were adults, whereas repeat sampled wild mice included animals classed as both adult and juvenile at the first sampling point. As such, we also repeated this analysis excluding all juveniles for comparability. Second, we used Levene’s test to ask whether the rate of epigenetic ageing differed according to (1) age category in wild mice (binary assessment; juvenile vs sub-adult/adult) or (2) source (lab vs wild).

### Ethical statement

Wild mouse work was conducted under Home Office license PPL PB0178858 held at the University of Oxford, and with research permits from the Islands Conservation Advisory Committee (ICAC), and Natural Resources Wales.

## Results

### Measurement of CpG site methylation levels from faecal samples

We used non-invasively collected faecal samples from laboratory and wild house mice (*Mus musculus* and *Mus musculus domesticus,* respectively) as a source of host DNA for the measurement of methylation levels at specific CpG sites. Sufficient amount of host DNA was extracted and subsequently sequenced despite use of a microbial DNA purification kit: methylation levels were successfully measured across all 73 CpG sites from the targeted house mouse genes (*Hsf4*, *Gm9312*, *Kcns1, Gm7325,* and *Prima1*; Table S2) in all samples from laboratory mice (*n*=80) and in 81% (183 out of 226) samples from wild mice (mean read depth per gene ranged 5,703–16,849 across samples). Methylation levels were successfully measured in 96% (217 out of 226) wild mouse samples when considering only those CpG sites included in the clock subsequently used to predict age in wild mice (11 CpG sites from genes *Hsf4* and *Kcns1,* see below). Repeatability of the method was assessed with 11 wild mouse DNA extractions processed twice (see Methods). Repeatability of methylation levels was 0.674 (standard error (SE)=0.052, *p*<0.001), while repeatability of epigenetic age estimates was 0.929 (SE=0.072, *p*<0.001).

### Construction of a non-invasive epigenetic clock

We first built an epigenetic clock using samples from C57BL/6 laboratory mice (*n*=50, one sample per animal) to generate a targeted epigenetic clock using methylation levels from 73 CpG sites across five genes that were previously associated with age in laboratory mice (*Hsf4*, *Gm9312*, *Kcns1, Gm7325,* and *Prima1;* Table. S1; Han et al., 2018). Elastic net regression identified 22 CpG sites from the five targeted genes that exhibited variability in methylation patterns with age; three from *Hsf4*, six from *Gm9312*, six from *Kcns1,* one from *Gm7325* and one from *Prima1* (Table S3). This epigenetic clock had a mean absolute error (MAE) of 7 days (∼0.7% of expected C57BL/6 lifespan; Schultz et al., 2020) in the training set (Pearson’s *r*=0.996, *p*<0.001; Fig. 1A). We validated the clock by applying it to an independent set of C57BL/6 mice that were not used in training the clock (*n*=15). Among these lab mice (one sample per animal), epigenetic age was also strongly correlated with chronological age (Pearson’s *r*=0.935, *p*<0.001, MAE=24 days; Fig. 1B). Neither sex nor sequencing run had a significant effect on epigenetic age (linear mixed effects model with two samples per animal: chronological age *F*=111.526, *p*<0.001; sex *F=*0.046, *p*=0.833; sequencing run *F=*1.602*, p*=0.217; animal ID included as a random effect). This demonstrates that non-invasive faecal samples can be used to generate an epigenetic clock in laboratory mice with equivalent or higher accuracy in estimating chronological age compared to a clock previously derived using blood samples (Han et al., 2018; MAE=35–41 days in two validation datasets).

As our aim was to develop a lab-based epigenetic clock that could be applied to wild mice, we built a second clock that did not include CpG sites that either showed non-parallel methylation level patterns across the two systems or for which we failed to acquire methylation levels in a substantial number of wild mouse samples (see Methods for more detail). For this clock, elastic net regression identified 11 CpG sites from genes *Hsp4* and *Kcns1* (Table S4), 7 of which were also included in the first clock (Table S3). Here, the slope deviated more from 1 (where 1 would indicate perfect positive linear relationship between chronological and epigenetic age) than did the first clock (slope estimate 0.756 ± 0.024 standard error vs 0.976 ± 0.013 standard error in training set; Fig 1A, 1C). However, the clock still had a high accuracy with a MAE of 23 days in the training set (Pearson’s *r*=0.977, *p*<0.001; Fig. 1C) and 26 days in the validation set (Pearson’s *r*=0.938, *p*<0.001; Fig. 1D). This second clock was then used for further analyses in the present study.

### Chronological age prediction in wild mice

We next applied the second lab-mouse derived epigenetic clock (Fig. 1C, 1D) to 201 faecal samples from 118 wild house mice to test if it can be used to estimate chronological age in wild individuals of unknown age. Mice of all available body sizes were included with the aim of capturing as much age variation as possible (body mass range 5.9–43.0g, mean 18.8, median 19.2). The epigenetic age of wild mice varied from −18 to 426 days (mean 213, median 193; 2 out of 201 samples (1%) had a negative epigenetic age).

Epigenetic age varied significantly between mice from different age categories that were assigned in the field using external characteristics (juvenile/sub-adult/adult; ANOVA, *F*_2,191_*=*131, *p*<0.001; Fig. 2A). Moreover, among 35 wild mice sampled twice between 30 and 340 days apart (mean 127, median 79), a great majority (*n=*30; 86%) were epigenetically older at the latter timepoint (one-tailed binomial test for H_0_ p=0.5, *p*<0.001). In general, the number of days between sampling time-points positively predicted change in epigenetic age (linear model, *F*_1,28_*=*7.814, *p=*0.009), but there was wide variation in the slope observed among individuals (Fig. 2B). Rate of epigenetic ageing was not predicted by any investigated variables (linear model with log-transformed response variable; trapping area *F*_1,23_*=*1.878, *p=*0.184; season *F*_2,23_*=*2.307, *p=*0.122; sex *F*_1,23_*=*0.237, *p=*0.631; reproductive activity *F*_1,23_*=*1.790, *p=*0.194 and body mass *F*_1,23_*=*0.013, *p*=0.909 at first sampling point). Together these results indicate that an epigenetic clock trained with samples from inbred lab mice can be used to provide an estimate of chronological age in outbred wild mice, though not one that is highly accurate.

**Figure 2.**
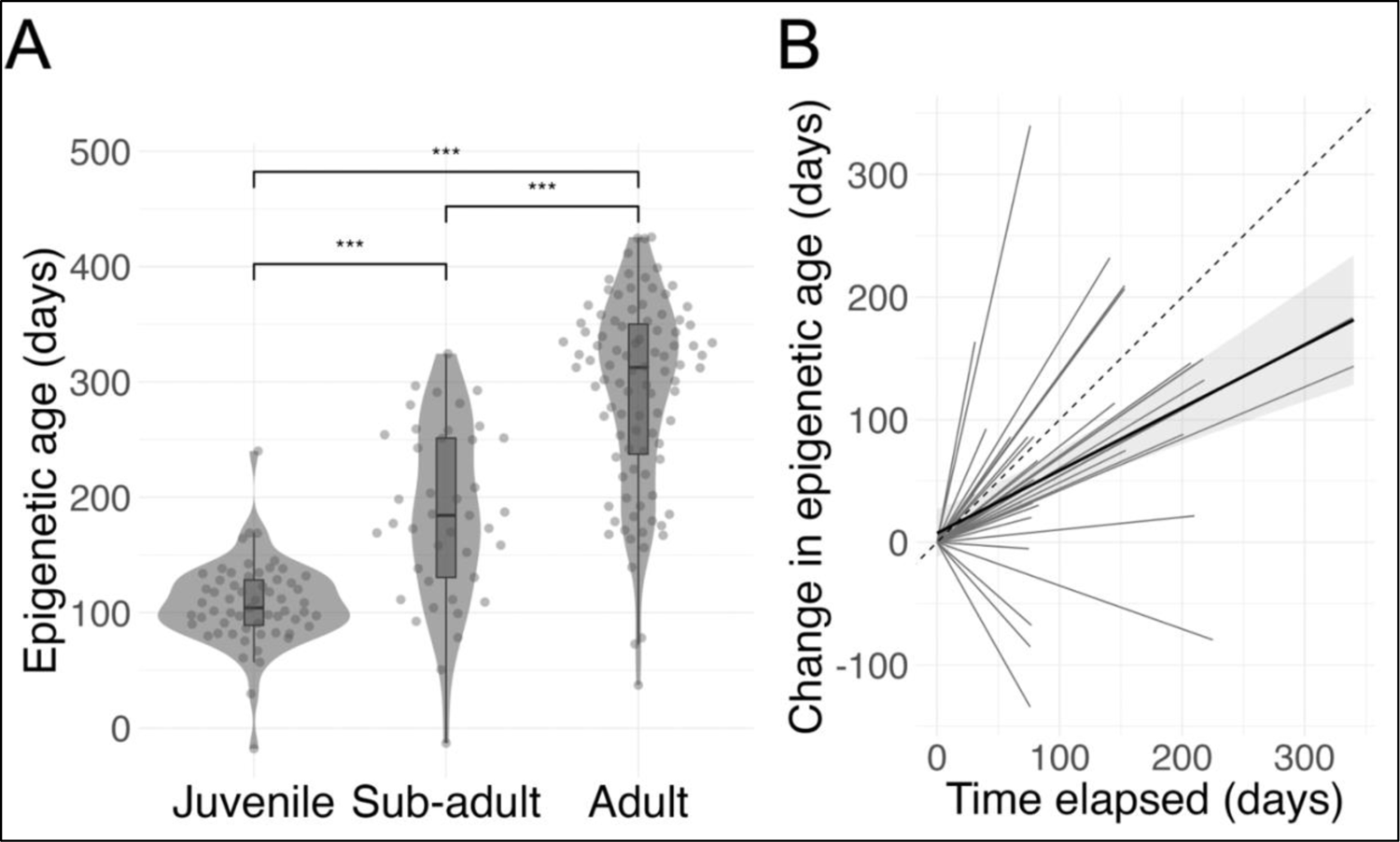
(**A**) Epigenetic age of wild mice phenotypically characterised as juvenile (*n*=62), sub-adult (*n*=37) or adult (*n*=95) predicted with an epigenetic clock built using C57BL/6 lab mice. Age category was assigned in the field using body size and appearance (see Methods). Median epigenetic age was 109 days in juveniles, 179 days in sub-adults and 296 days in adults. Statistical differences between different age categories were tested with Wilcoxon rank sum tests (***; *p*<0.001). (**B**) Change in epigenetic age between two timepoints in wild mice sampled twice between 30 and 340 days apart (*n*=35). Grey lines represent individual mice, black line is a linear regression line with 95% confidence interval bands and dashed line is a reference line of *y=x* (the hypothetical relationship if chronological age and epigenetic age estimates were exactly equivalent). Epigenetic age increased with time for 30 out of 35 (86%) mice.

### Multiple times higher methylation levels in wild compared to lab mice

To investigate whether wild mice had higher methylation levels than lab mice (which would lead to a higher epigenetic age estimate) in early life and beyond, we explored the relationship between methylation levels and body mass across mice of all sizes. Methylation levels were strongly predicted by source (brm model; source, posterior mean 1.10, 95% credible intervals (CI) 0.90–1.31; body mass, posterior mean 0.05, 95% credible intervals (CI) 0.05–0.06; model includes Animal ID random effect as well as a nested random effect Gene/Position), such that wild mice had higher levels of methylation in the genes showing parallel trends across lab and wild mice (Fig. S1–4).

We next assessed whether this higher methylation level in wild mice resulted in higher epigenetic age for a given chronological age in wild compared to laboratory mice. In the absence of known chronological age for wild mice, we used body mass to provide an upper limit age estimate for individuals classed as juveniles. Others have reported that 12–13-day-old wild house mice from mainland Europe weigh around 7g (range 3.6–10.5g, mean 6.8; Gerber et al., 2021) and another study showed that 14 day old wild-derived but captive house mice from Gough Island (home to the largest wild house mice recorded) weigh around 8.5g (range ∼7–10.5g, raw data not available; Gray et al., 2015; Fig. S5). Thus, irrespective of context, house mice between 12–14 days typically are expected to weigh 7–10.5g (Gerber et al., 2021; Gray et al., 2015). We therefore examined epigenetic age from a random cross-sectional set of juvenile wild Skokholm Island mice that fall within this body mass range and thus which we expect to be no more than 25 days old (*n*=19, body mass 6.1–10.4g with a mean of 8.5g). Among these individuals, the mean epigenetic age estimate was 106 days (range −18–169, median 104; 1 sample had a negative epigenetic age; for comparison, the epigenetic age in lab mice <14d of age (*n*=7) ranged 22–34 with a mean of 29 days), which is more than four times their expected chronological age (Fig. S5). Looking across wild mice of all body masses, the same pattern was maintained, with wild mice having higher epigenetic age estimates than lab mice across all body sizes (generalized additive model; body mass, *F*=38.10, *p*<0.001, body mass*source, *F*=13.40, *p*<0.001; Fig. 3A).

**Figure 3.**
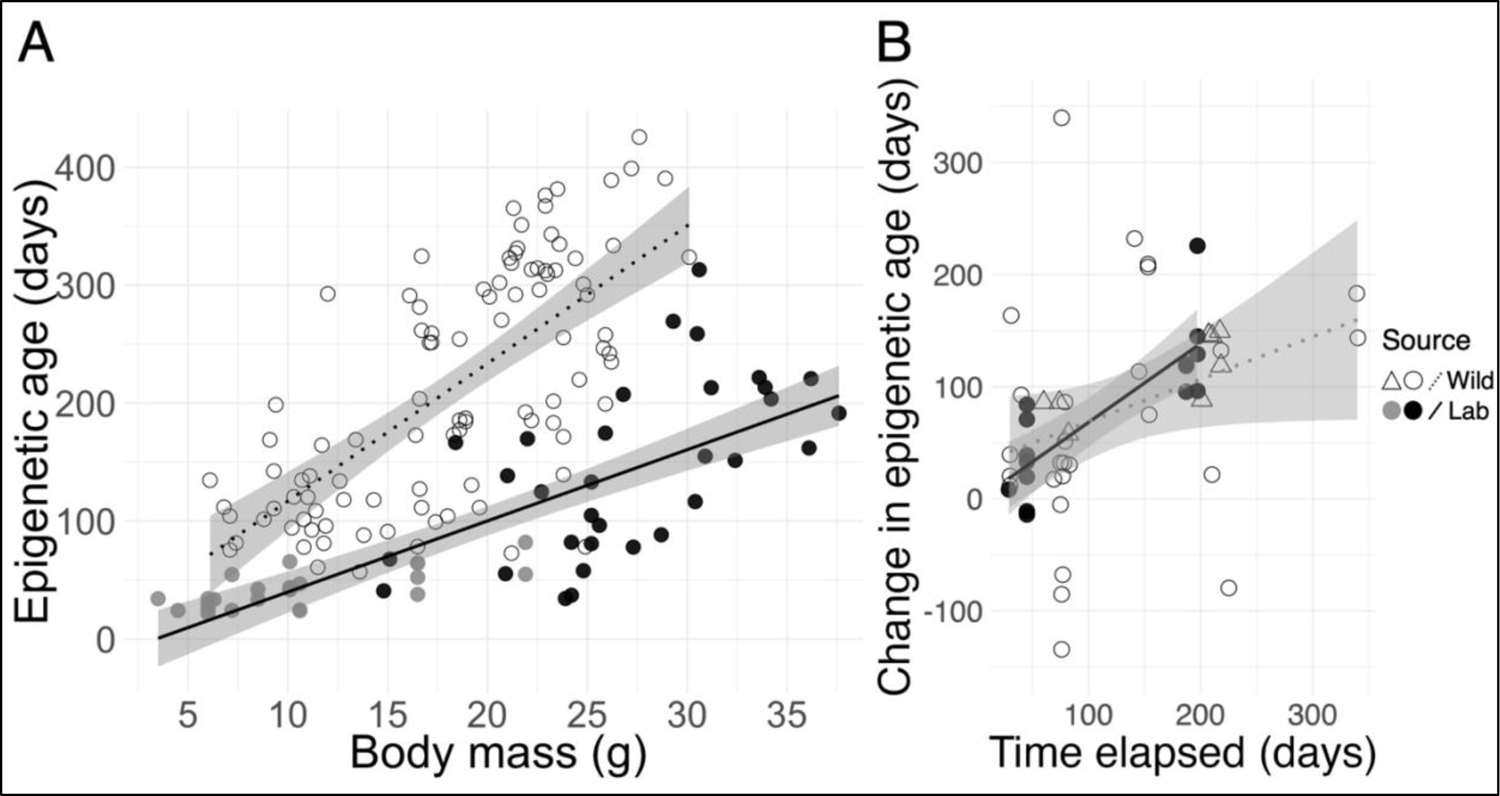
Wild mice have elevated epigenetic age compared to lab mice for a given body mass, but do not show a different rate of epigenetic ageing in adulthood. (**A**) Epigenetic age of wild (*n*=99) and laboratory mice (*n*=57) in relation to body mass. Empty circles are wild mice and filled circles are lab mice; black circles are lab mice for which body mass was recorded during sample collection, and grey circles are lab mice for which body mass was estimated post hoc (see Methods). (**B**) Change in epigenetic age in relation to time elapsed in laboratory (*n*=15) and wild mice (*n*=35) sampled twice a minimum 26 days apart. Body mass varies 19.2–26.3g for lab mice and 7.0–37.7g for wild mice. Empty triangles (*n*=8) are wild mice classed as ‘juvenile’ at first timepoint, empty circles (*n*=27) are wild mice classed as ‘sub-adult’ or ‘adult’ at first timepoint and filled circles are lab mice. Lines are linear regression lines (*dashed* = wild mice, *solid* = lab mice) and shading indicates 95% confidence intervals.

To further examine whether older epigenetic age profiles among wild mice might be due to accelerated ageing through exposure to environmental stressors, such as food shortage or climatic variation, we studied the rate of epigenetic ageing across lab and wild mice for which two timepoints were available (lab *n=*15; wild *n*=35). While the rate of epigenetic ageing appeared slightly shallower in wild mice (Fig. 3B), source did not predict rate of epigenetic ageing (linear model: source, *F*_1,39_=0.010, *p*=0.923; controlling for sex, *F*_1,39_=0.408, *p*=0.527, and body mass at first timepoint, *F*_1,39_=0.105, *p*=0.747). Lab mice included in this analysis were all adults, while the longitudinal data of wild mice included adult and juvenile mice; however, exclusion of juvenile mice (≤15g of body mass and reproductively inactive, *n*=8) did not appear to have a strong effect on these trends (linear model: source, *F*_1,31_=0.004, *p*=0.948; controlling for sex, *F*_1,31_=0.213, *p*=0.648, and body mass at first timepoint, *F*_1,31_=0.226, *p*=0.638). Further, although the variance in estimates of epigenetic ageing rates was lower for juvenile wild mice than for non-juvenile (sub-adult/adult) wild mice, this variance difference was not statistically significant (Levene’s test, *F*_1,33_=1.770, *p*=0.193; Pearson’s correlation between change in epigenetic age and days elapsed: juvenile wild mice, *r*=0.79, *p*=0.021; non-juvenile wild mice, *r*=0.32, *p*=0.102; Fig. 3B). Similarly, higher variation in epigenetic ageing rates observed in wild compared to lab mice was not statistically significant (Levene’s test, *F*_1,48_=3.428, *p*=0.070; Pearson’s correlation between change in epigenetic age and days elapsed: lab mice, *r*=0.83, *p*<0.001; wild mice, *r*=0.36, *p*=0.033; Fig. 3B).

## Discussion

Here, we tested an approach for estimating age in wild house mice, by building an epigenetic clock using samples from inbred C57BL/6 laboratory mice and using it to estimate age in outbred wild mice of unknown chronological age. Faecal samples were used as a source of host DNA and proved suitable for measuring of DNA methylation and epigenetic age, indicating their potential as a non-invasive alternative to the blood or tissue samples more commonly used in epigenetic clocks (Han et al., 2018). The clock effectively distinguished wild juveniles from adults, and typically showed increases in predicted age over time among repeat-captured individuals. The success rate of the latter (86% individuals predicted older at a later time point) was similar to what has been previously been detected in a wild baboon study (Anderson et al., 2021).

However, while the clock accurately predicted age in an independent set of laboratory mice (with error of ± 26 days; ∼3% of expected C57BL/6 lifespan), we observed high variation among wild mice in how their epigenetic age changed over chronological time, suggesting our clock had far less accuracy in predicting chronological age in this different ecological context. Others have had better success in applying clocks built with captive individuals to wild individuals (e.g., Mayne et al., 2022 in green turtles; Robeck et al., 2021 in cetaceans; correlation between chronological and epigenetic age in these studies 0.67–0.98 *vs* 0.36 between change in time and change in epigenetic age in our study). However, these studies have built epigenetic clocks using samples from captive individuals where there is genetic and environmental variation, such as animals from zoos or outdoor enclosures. Our clock was built with samples from inbred lab mice housed under very stable environmental conditions, but applied to wild mice that are outbred and exposed to a highly variable temperate climate. Studies of different lab strains have confirmed that epigenetic clocks may behave differently in different genetic backgrounds. For instance, Han et al demonstrated DBA/2 mice to be up to twice as old epigenetically as C57BL/6 mice (Han et al., 2018). Moreover, DNA methylation may be influenced by inbreeding (Han et al., 2021; Venney et al., 2016) and environmental factors (Zocher et al., 2021; Parrott et al., 2014; Viitaniemi et al., 2019), and wild animals are generally exposed to more variable environments than their inbred lab counterparts. As such, the contrasting genetic and environmental backgrounds in our mouse systems may partly explain why age estimates in wild mice based on a clock from lab mice had low accuracy.

Technical factors may have also contributed to the low accuracy of a lab animal-based clock when used to estimate age in wild animals. The accuracy of chronological age prediction particularly for older mice might have been affected by the relatively lower number of lab mice aged over 3 months in the training set. This is because the clock could be more inclined to capture patterns prevalent in younger mice, potentially resulting in an incomplete representation of the diverse epigenetic changes associated with ageing later in life. Further, while all samples from lab mice were preserved immediately after defecation, the time between defecation and sample preservation varied in wild mice (where samples were collected from traps which animals had been in overnight, up to 13 hours). It is possible some DNA degradation occurred before the samples were preserved in a stabilising buffer, affecting the methylation profiles. It is also possible that host cell profiles vary to some extent between faecal samples from lab and wild mice. Since methylation levels vary between tissue types (Han et al., 2018), differences in faecal cell profiles could have contributed to our findings on methylation levels in lab vs wild mice.

As our study is the first to develop a lab-based epigenetic clock based on faecal samples and apply it to a wild setting, we cannot assess whether the relatively low accuracy in chronological ageing we achieved is species- or sample type-specific. It is also possible that epigenetic changes with age are comparable across settings (lab and wild) but that biological ageing varies more in the wild than it does in the lab, obscuring any chronological signal in epigenetic markers. Further studies would be needed to understand whether this is the case, and whether lab-based clocks might still hold value for estimating either chronological or biological ageing in wild animals. An alternative clock to ours could be trained using samples from captive individuals with greater genetic diversity, such as more genetically diverse lab mice rather than an inbred strain, or using samples from free-living populations where chronological age can be accurately estimated, e.g., outbred semi-natural populations where individuals can be tracked from birth (Gerber et al., 2021).

The epigenetic age of wild mice from Skokholm Island varied from −18 to 426 days (mean 213, median 193; 2 (1% of all 201) samples had negative epigenetic ages, −18 and −13 days). The presence of negative predicted ages in wild individuals may be due to measurement error, but it may also be due to the inherent biological differences in age-related epigenetic changes in wild vs lab populations. DNA methylation-based age predictions may reflect the distinct age trajectories and relative age of wild individuals compared to their lab counterparts, highlighting the importance of considering the natural variation in epigenetic ageing processes across different populations.

Despite our epigenetic clock very accurately predicting the chronological age of laboratory mice, several lines of evidence suggest that in wild mice, our epigenetic age estimates were overestimates of chronological age. First, among juveniles, for which body mass is an accurate predictor of age across both wild and lab mice (Jax, 2022a; Jax, 2022b; Gerber et al., 2021; Gray et al., 2015), epigenetic age estimates were several times higher than their expected chronological age from body mass (Gerber et al., 2021; Gray et al., 2015). Second, we found that wild mice exhibited higher levels of CpG site methylation (and subsequently several times higher estimates of epigenetic age) across all body masses, compared to lab mice. While accelerated weight gain in *ad libitum*-fed lab mice may contribute to lower methylation levels among adult lab mice compared to wild mice, it may also be that a more challenging environment experienced by wild mice increases methylation and consequently accelerates epigenetic clocks. Our comparison of methylation levels and epigenetic age of wild vs lab mice is specific to a comparison with the C57BL/6 strain. However, it is perhaps noteworthy that the approximately 5–10-fold difference in epigenetic age between C57BL/6 lab mice and wild mice found here is larger than the previously reported 2-fold difference in epigenetic age between C57BL/6 mice and another inbred lab strain, DBA/2 (Han et al., 2018).

To test whether the older epigenetic age profile of wild mice could be explained by accelerated ageing post-weaning (i.e., from when they are trappable) we investigated the rate of epigenetic ageing using individuals captured and sampled twice over time. If anything, the rate of epigenetic aging appeared slightly slower and more variable in wild mice, though this observation relied on a small sample size and was not statistically significant. Various factors can influence methylation levels and these factors could differ between lab and wild mice settings here, such as abiotic factors (e.g., temperature), inbreeding (Han et al., 2021; Venney et al., 2016), and food shortage (as caloric restriction may slow epigenetic ageing, Maegawa et al., 2017; Hahn et al., 2017). Moreover, as we observed heightened epigenetic age in wild compared to lab mice even during the first ∼2 weeks of life, we suggest that peri- and early postnatal effects on offspring DNA methylation may vary between laboratory and wild mice. Various human, mouse, and other animal studies have demonstrated the association between prenatal maternal experience (such as food shortage, diet, infection, substance exposure, and stress) and offspring DNA methylation patterns, with differences from the prenatal (foetal) phase still detectable in later life (Tobi et al., 2009; Heijmans et al., 2008; Lan et al., 2013; Richetto et al., 2017; Camerota et al., 2021; Joubert et al., 2016; Kertes et al., 2016; Vangeel et al., 2017).

In our present study we used targeted sequencing to measure methylation levels in genes of interest. As such, while methylation levels are higher in wild than lab mice in these targeted genes, this may not be the case for genome-wide methylation. Further, when comparing methylation levels and epigenetic age estimates across lab and wild mice, we have used body mass as a proxy for age across lab and wild mice. Body mass is, however, only a rough proxy and mass-age relationships are likely to vary somewhat across lab and wild. To more definitely explore whether methylation levels and biological age for a given chronological age vary across these contexts, comparisons of epigenetic aging patterns in (semi-)wild mice of known chronological age with these lab and wild populations would be very valuable.

While our approach of training an epigenetic clock with lab individuals and using it to estimate age in wild individuals did not allow accurate estimation of chronological age, our results demonstrate such an approach can still be effective in distinguishing between juvenile and adult individuals. Such information may be useful in contexts where a faecal deposit is found but the individual is not observed, such as in field-based projects of animals that are hard or impossible to capture. At the same time, this method can provide interesting insights into biological ageing when applied to wild animals of known chronological age or to individuals sampled longitudinally such that changes in epigenetic age can be estimated (De Paoli-Iseppi et al., 2017; Powell & Proulx, 2003; Brivio et al., 2015). Considering the much greater variability in epigenetic ageing rates we observed in wild compared to laboratory animals, our results suggest wild systems may provide an informative environment in which to study drivers of epigenetic age acceleration. Our current study did not have sufficient power to ask why wild individuals might vary so widely in epigenetic aging rates, and most longitudinally sampled individuals were adults. Further work using a larger sample size and greater coverage of different life stages would be valuable to systematically explore potential drivers of this fascinating variation.

In summary, our data indicate the potential to use a non-invasive, DNA methylation-based epigenetic clock built with samples from laboratory mice to estimate the age of wild mice. While this approach did not provide highly accurate estimates of chronological age, it can be used to measure variation in biological ageing in future longitudinal studies, making it a promising tool for studies of ontogeny and senescence in wild settings.

## Acknowledgements

We thank Giselle Eagle and Richard Brown, the wardens of Skokholm Island, the Friends of Skokholm and Skomer, the Wildlife Trust of South and West Wales and field assistants for their help in enabling the wild mouse data collection.

## Author information

### Contributions

EH and SCLK set up the wild mouse study system. EH, SJ, AR and KW collected the samples. AO and TJL developed the software Apollo. EH conducted the laboratory work and analysed the data with support from TJL, SCLK and AR. EH wrote the manuscript with contributions from all authors.

### Data Accessibility

Data and R scripts are available at https://github.com/eveliinahanski/musmus_epiage

## References

Ambatipudi S, Horvath S, Perrier F, et al. DNA methylome analysis identifies accelerated epigenetic ageing associated with postmenopausal breast cancer susceptibility. Eur J Cancer. 2017;75:299–307. doi:10.1016/j.ejca.2017.01.014

Anderson JA, Johnston RA, Lea AJ, et al. High social status males experience accelerated epigenetic ageing in wild baboons. Elife. 2021;10:e66128. Published 2021 Apr 6. doi:10.7554/eLife.66128

Bates D, Mächler M, Bolker B, Walker S (2015). “Fitting Linear Mixed-Effects Models Using lme4.” Journal of Statistical Software, 67(1), 1–48. doi:10.18637/jss.v067.i01.

Berry RJ, Jakobson ME. Life and death in an island population of the house mouse. Exp Gerontol. 1971;6: 187–197.

Bors EK, Baker CS, Wade PR, et al. An epigenetic clock to estimate the age of living beluga whales. Evol Appl. 2021;14(5):1263–1273. Published 2021 Feb 3. doi:10.1111/eva.13195

Brivio F, Grignolio S, Sica N, Cerise S, Bassano B. Assessing the Impact of Capture on Wild Animals: The Case Study of Chemical Immobilisation on Alpine Ibex. PLoS One. 2015;10(6):e0130957. Published 2015 Jun 25. doi:10.1371/journal.pone.0130957

Paul-Christian Bürkner (2017). brms: An R Package for Bayesian Multilevel Models Using Stan. Journal of Statistical Software, 80(1), 1–28. doi:10.18637/jss.v080.i01

Camerota M, Graw S, Everson TM, et al. Prenatal risk factors and neonatal DNA methylation in very preterm infants. Clin Epigenetics. 2021;13(1):171. Published 2021 Sep 10. doi:10.1186/s13148-021-01164-9

Cao X, Li W, Wang T, et al. Accelerated biological ageing in COVID-19 patients. Nat Commun. 2022;13(1):2135. Published 2022 Apr 19. doi:10.1038/s41467-022-29801-8

De Paoli-Iseppi R, Deagle BE, McMahon CR, Hindell MA, Dickinson JL, Jarman SN. Measuring Animal Age with DNA Methylation: From Humans to Wild Animals. Front Genet. 2017;8:106. Published 2017 Aug 17. doi:10.3389/fgene.2017.00106

Fairfield EA, Richardson DS, Daniels CL, Butler CL, Bell E, Taylor MI. Ageing European lobsters (*Homarus gammarus*) using DNA methylation of evolutionarily conserved ribosomal DNA. Evol Appl. 2021;14(9):2305–2318. Published 2021 Sep 23. doi:10.1111/eva.13296

Ferrari, Manuela; Lindholm, Anna K; König, Barbara (2015). The risk of exploitation during communal nursing in house mice, *Mus musculus domesticus*. Animal Behaviour, 110:133–143.

Flurkey, Currer, and Harrison, 2007. ’The mouse in biomedical research.’ in James G. Fox (ed.), American College of Laboratory Animal Medicine series (Elsevier, AP: Amsterdam; Boston).

Friedman J, Hastie T, Tibshirani R. Regularization Paths for Generalized Linear Models via Coordinate Descent. J Stat Softw. 2010;33(1):1–22.

Gardner ST, Bertucci EM, Sutton R, Horcher A, Aubrey D, Parrott BB. Development of DNA methylation-based epigenetic age predictors in loblolly pine (Pinus taeda). Mol Ecol Resour. 2023;23(1):131–144. doi:10.1111/1755-0998.13698

Georgountzou, A., & Papadopoulos, N. G. (2017). Postnatal Innate Immune Development: From Birth to Adulthood. Frontiers in immunology, 8, 957.10.3389/fimmu.2017.00957

Gerber N, Auclair Y, König B, Lindholm AK. Population density and temperature influence the return on maternal investment in wild house mice. Front Ecol Evol. 2021;8:602359. 10.3389/fevo.2020.602359

Gray MM, Parmenter MD, Hogan CA, et al. Genetics of Rapid and Extreme Size Evolution in Island Mice. Genetics. 2015;201(1):213–228. doi:10.1534/genetics.115.177790

Han, T., Wang, F., Song, Q., Ye, W., Liu, T., Wang, L., & Chen, Z. J. (2021). An epigenetic basis of inbreeding depression in maize. Science advances, 7(35), eabg5442. 10.1126/sciadv.abg5442

Han Y, Eipel M, Franzen J, et al. Epigenetic age-predictor for mice based on three CpG sites. Elife. 2018;7:e37462. Published 2018 Aug 24. doi:10.7554/eLife.37462

Hahn, O., Grönke, S., Stubbs, T. M., Ficz, G., Hendrich, O., Krueger, F., Andrews, S., Zhang, Q., Wakelam, M. J., Beyer, A., Reik, W., & Partridge, L. (2017). Dietary restriction protects from age-associated DNA methylation and induces epigenetic reprogramming of lipid metabolism. Genome biology, 18(1), 56. 10.1186/s13059-017-1187-1

Harvanek ZM, Fogelman N, Xu K, Sinha R. Psychological and biological resilience modulates the effects of stress on epigenetic ageing. Transl Psychiatry. 2021;11(1):601. Published 2021 Nov 27. doi:10.1038/s41398-021-01735-7

Heijmans BT, Tobi EW, Stein AD, et al. Persistent epigenetic differences associated with prenatal exposure to famine in humans. Proc Natl Acad Sci U S A. 2008;105(44):17046–17049. doi:10.1073/pnas.0806560105

Ito H, Udono T, Hirata S, Inoue-Murayama M. Estimation of chimpanzee age based on DNA methylation. Sci Rep. 2018;8(1):9998. Published 2018 Jul 3. doi:10.1038/s41598-018-28318-9

Jarman SN, Polanowski AM, Faux CE, et al. Molecular biomarkers for chronological age in animal ecology. Mol Ecol. 2015;24(19):4826–4847. doi:10.1111/mec.13357

JAX^®^. Body weight information for aged C57BL/6J. Available at: https://www.jax.org/jax-mice-and-services/strain-data-sheet-pages/body-weight-chart-aged-b6 (Accessed: 30.11.2022)

JAX^®^. Body weight information for C57BL/6J. Available at: https://www.jax.org/jax-mice-and-services/strain-data-sheet-pages/body-weight-chart-000664 (Accessed: 30.11.2022)

Joubert BR, Felix JF, Yousefi P, et al. DNA Methylation in Newborns and Maternal Smoking in Pregnancy: Genome-wide Consortium Meta-analysis. Am J Hum Genet. 2016;98(4):680–696. doi:10.1016/j.ajhg.2016.02.019

Joyce BT, Gao T, Zheng Y, et al. Epigenetic Age Acceleration Reflects Long-Term Cardiovascular Health. Circ Res. 2021;129(8):770–781. doi:10.1161/CIRCRESAHA.121.318965

Kerepesi C, Meer MV, Ablaeva J, et al. Epigenetic ageing of the demographically non-ageing naked mole-rat. Nat Commun. 2022;13(1):355. Published 2022 Jan 17. doi:10.1038/s41467-022-27959-9

Kertes DA, Kamin HS, Hughes DA, Rodney NC, Bhatt S, Mulligan CJ. Prenatal Maternal Stress Predicts Methylation of Genes Regulating the Hypothalamic-Pituitary-Adrenocortical System in Mothers and Newborns in the Democratic Republic of Congo. Child Dev. 2016;87(1):61–72. doi:10.1111/cdev.12487

Lan X, Cretney EC, Kropp J, et al. Maternal Diet during Pregnancy Induces Gene Expression and DNA Methylation Changes in Fetal Tissues in Sheep. Front Genet. 2013;4:49. Published 2013 Apr 5. doi:10.3389/fgene.2013.00049

Larison B, Pinho GM, Haghani A, et al. Epigenetic models developed for plains zebras predict age in domestic horses and endangered equids. Commun Biol. 2021;4(1):1412. Published 2021 Dec 17. doi:10.1038/s42003-021-02935-z

Lemaître JF, Rey B, Gaillard JM, et al. DNA methylation as a tool to explore ageing in wild roe deer populations. Mol Ecol Resour. 2022;22(3):1002–1015. doi:10.1111/1755-0998.13533

Maegawa, S., Lu, Y., Tahara, T., Lee, J. T., Madzo, J., Liang, S., Jelinek, J., Colman, R. J., & Issa, J. J. (2017). Caloric restriction delays age-related methylation drift. Nature communications, 8(1), 539. 10.1038/s41467-017-00607-3

Mayne B, Mustin W, Baboolal V, et al. Age prediction of green turtles with an epigenetic clock. Mol Ecol Resour. 2022;22(6):2275–2284. doi:10.1111/1755-0998.13621

Moore LD, Le T, Fan G. DNA methylation and its basic function. Neuropsychopharmacology. 2013;38(1):23–38. doi:10.1038/npp.2012.112

Morales Berstein F, McCartney DL, Lu AT, et al. Assessing the causal role of epigenetic clocks in the development of multiple cancers: a Mendelian randomization study. Elife. 2022;11:e75374. Published 2022 Mar 29. doi:10.7554/eLife.75374

Parrott BB, Bowden JA, Kohno S, et al. Influence of tissue, age, and environmental quality on DNA methylation in Alligator mississippiensis. Reproduction. 2014;147(4):503–513. Published 2014 Mar 2. doi:10.1530/REP-13-0498

Peng C, Cardenas A, Rifas-Shiman SL, et al. Epigenetic age acceleration is associated with allergy and asthma in children in Project Viva. J Allergy Clin Immunol. 2019;143(6):2263–2270.e14. doi:10.1016/j.jaci.2019.01.034

Petkovich DA, Podolskiy DI, Lobanov AV, Lee SG, Miller RA, Gladyshev VN. Using DNA Methylation Profiling to Evaluate Biological Age and Longevity Interventions. Cell Metab. 2017;25(4):954–960.e6. doi:10.1016/j.cmet.2017.03.016

Pinho GM, Martin JGA, Farrell C, et al. Hibernation slows epigenetic ageing in yellow-bellied marmots. Nat Ecol Evol. 2022;6(4):418–426. doi:10.1038/s41559-022-01679-1

Polanowski AM, Robbins J, Chandler D, Jarman SN. Epigenetic estimation of age in humpback whales. Mol Ecol Resour. 2014;14(5):976–987. doi:10.1111/1755-0998.12247

Powell RA, Proulx G. Trapping and marking terrestrial mammals for research: integrating ethics, performance criteria, techniques, and common sense. ILAR J. 2003;44(4):259–276. doi:10.1093/ilar.44.4.259

Prado NA, Brown JL, Zoller JA, et al. Epigenetic clock and methylation studies in elephants. ageing Cell. 2021;20(7):e13414. doi:10.1111/acel.13414

R Core Team (2023). _R: A Language and Environment for Statistical Computing_. R Foundation for Statistical Computing, Vienna, Austria. <https://www.R-project.org/>.

Richetto J, Massart R, Weber-Stadlbauer U, Szyf M, Riva MA, Meyer U. Genome-wide DNA Methylation Changes in a Mouse Model of Infection-Mediated Neurodevelopmental Disorders. Biol Psychiatry. 2017;81(3):265–276. doi:10.1016/j.biopsych.2016.08.010

Robeck TR, Fei Z, Lu AT, et al. Multi-species and multi-tissue methylation clocks for age estimation in toothed whales and dolphins. Commun Biol. 2021;4(1):642. Published 2021 May 31. doi:10.1038/s42003-021-02179-x

Schultz, M. B., Kane, A. E., Mitchell, S. J., MacArthur, M. R., Warner, E., Vogel, D. S., Mitchell, J. R., Howlett, S. E., Bonkowski, M. S., & Sinclair, D. A. (2020). Age and life expectancy clocks based on machine learning analysis of mouse frailty. Nature communications, 11(1), 4618. 10.1038/s41467-020-18446-0

Spangenberg E, Wallenbeck A, Eklöf AC, Carlstedt-Duke J, Tjäder S. Housing breeding mice in three different IVC systems: maternal performance and pup development. Lab Anim. 2014;48(3):193–206. doi:10.1177/0023677214531569

Stoffel, M. A., Nakagawa, S. and Schielzeth, H. (2017), rptR: repeatability estimation and variance decomposition by generalized linear mixed-effects models. Methods Ecol Evol, 8: 1639???1644. doi:10.1111/2041-210X.12797

Stubbs TM, Bonder MJ, Stark AK, et al. Multi-tissue DNA methylation age predictor in mouse. Genome Biol. 2017;18(1):68. Published 2017 Apr 11. doi:10.1186/s13059-017-1203-5

Sullivan IR, Adams DM, Greville LJS, Faure PA, Wilkinson GS. Big brown bats experience slower epigenetic ageing during hibernation. Proc Biol Sci. 2022;289(1980):20220635. doi:10.1098/rspb.2022.0635

Tangili M, Slettenhaar AJ, Sudyka J, et al. DNA methylation markers of age(ing) in non-model animals. Mol Ecol. 2023;32(17):4725–4741. doi:10.1111/mec.17065

Thompson MJ, vonHoldt B, Horvath S, Pellegrini M. An epigenetic ageing clock for dogs and wolves. ageing (Albany NY). 2017;9(3):1055–1068. doi:10.18632/ageing.101211

Tobi EW, Lumey LH, Talens RP, et al. DNA methylation differences after exposure to prenatal famine are common and timing- and sex-specific. Hum Mol Genet. 2009;18(21):4046–4053. doi:10.1093/hmg/ddp353

Vangeel EB, Pishva E, Hompes T, et al. Newborn genome-wide DNA methylation in association with pregnancy anxiety reveals a potential role for *GABBR1*. Clin Epigenetics. 2017;9:107. Published 2017 Oct 3. doi:10.1186/s13148-017-0408-5

Venney, C. J., Johansson, M. L., & Heath, D. D. (2016). Inbreeding effects on gene-specific DNA methylation among tissues of Chinook salmon. Molecular ecology, 25(18), 4521–4533. 10.1111/mec.13777

Viitaniemi HM, Verhagen I, Visser ME, Honkela A, van Oers K, Husby A. Seasonal Variation in Genome-Wide DNA Methylation Patterns and the Onset of Seasonal Timing of Reproduction in Great Tits. Genome Biol Evol. 2019;11(3):970–983. doi:10.1093/gbe/evz044

Wilkinson GS, Adams DM, Haghani A, et al. DNA methylation predicts age and provides insight into exceptional longevity of bats [published correction appears in Nat Commun. 2021 May 5;12(1):2652] [published correction appears in Nat Commun. 2022 Sep 7;13(1):5266]. *Nat Commun*. 2021;12(1):1615. Published 2021 Mar 12. doi:10.1038/s41467-021-21900-2

Wright PGR, Mathews F, Schofield H, et al. Application of a novel molecular method to age free-living wild Bechstein’s bats. Mol Ecol Resour. 2018;18(6):1374–1380. doi:10.1111/1755-0998.12925

Yousefzadeh, M., Henpita, C., Vyas, R., Soto-Palma, C., Robbins, P., & Niedernhofer, L. (2021). DNA damage-how and why we age?. eLife, 10, e62852. 10.7554/eLife.62852

Zocher S, Overall RW, Lesche M, Dahl A, Kempermann G. Environmental enrichment preserves a young DNA methylation landscape in the aged mouse hippocampus. Nat Commun. 2021;12(1):3892. Published 2021 Jun 23. doi:10.1038/s41467-021-23993-1

