## Supplementary Material for "Epigenetic age estimation of wild mice using faecal samples"

### Title

#### Supplementary material

**Supplementary Table 1 is provided as an Excel spreadsheet for easier inspection.**

**Supplementary Table 2.** Primers used to target five *Mus musculus* genes (*Hsf4, Gm9312, Fm7325*, *Kcns1*, *Prima1*), generated based on Han et al (2018).

| Gene | Primer sequence |
|---|---|
| *Hsf4* | GTGAGTAGTAAGGTGGGATAAATTGTAGAAAAAATG (Forward),<br>TCCCTACTCTCCTACACTCCTCTCAAAACTTA (Reverse) |
| *Gm9312* | TTGTTTTGGGGTATTAGAAATTTTTTT (Forward),<br>CCTAACCATACTAAACCAAATCTCTATATCTAAAT (Reverse) |
| *Gm7325* | TGTTGGTTGAGGATAAAGAGTAGATAGTTTAGTAGAGT (Forward),<br>TTCCCTTTACAAATACAAATCCTACCATA (Reverse) |
| *Kcns1* | GGTTGAGAGGGTGGTAGAAGAAGTTG (Forward),<br>ACTCCCCTCCATCCCTACCATATACATCCA (Reverse) |
| *Prima1* | TTGTGTTTAATTAGGAGAGGTAAATTATGAATTAGGTTTATA (Forward),<br>CAAAATTAATTACACCAACTTATAACCTACTATTC (Reverse) |

**Supplementary Table 3.** Elastic net regression used to build an epigenetic clock identified 22 CpG sites (in red) from five targeted genes (*Hsf4, Gm9312, Gm7325, Kcns1, Prima1*): three from *Hsf4*, six from *Gm9312*, one from *Gm7325*, eleven from *Kcns1* and one from *Prima1*.

| **Gene** | **Sequence** |
|---|---|
| *Hsf4* | GGAAGGTATTAATGTTGGTATTTTTGGTTTTGTTTATGTGTTTCGGATGGTGTTTTT<br>TGTTTGTAGGTATTTGCGTTGCGAGGCGATGATAGTCGATGGCGTTCGGAAGATT<br>TGAGTCGATTGTTGGGAGAGGTG |
| *Gm9312* | AGGTGTGGGCGTAGTCGGAGGGTATTGGGTATCGGGTATTAAGCGCGGAAGTTT<br>ATTAGGTGTTTAGGGCGTAGCGCGATTTTCGCGATTTTAGTTTTTCGTTCGTTGCG<br>GGTTACGTTATCGTTTATTTTTTCGATGGTCGTCGCGGTGTTTCGGTATTGGGTCG<br>AGCGCGTGGTGAAATTAGAGGTCGTGGGCGTTTTGTAGTTTTTAAGGGTTTCGGT<br>TACGA |
| *Gm7325* | TTTTTTTATGTTTTGGGAGTTTAGTCGGCGGGTTAGTCGGCGTAGTAAGGGTAGG<br>AGGTGTTGTCGGGTTAGGCGGGTAATAGGTAGTAGTAGGCGGGATAGTAGCGAT<br>CGAAGTATTATCGGGAGTAATGGAACGGG |
| *Kcns1* | CGCGTGTTGGGAGTTAGTAGTAGGCGCGACGATATTTCGAAGTTGAATTAAGCGA<br>TGTAGAAGTATTTTAGGCGGCGTAGTATCGGGTCGTCGCGTATTTTTTTGCGTTG<br>CGATTCGCGGTTATTGTAGTTATCGTCGTCGTCGTTTTTCGAGTTTGGTATTTCGG<br>TAGGTTG |
| *Prima1* | TTGTGTTTAATTAGGAGAGGTAAATTATGAATTAGGTTTATATTTTTCGGGTGGG<br>GGGTAACGACGGTTTTTTTTGGGTGAGGAAAAGATTATAGGGAGGTTTGAGTAA<br>GAGTCGGGAATAGTAGGTTATAAGTTGGTGTAATTAATTTTG |

**Supplementary Table 4**. Elastic net regression used to build an epigenetic clock identified 11 CpG sites (in red) from two of five targeted genes (*Hsf4, Gm9312, Gm7325, Kcns1, Prima1*): two from *Hsf4* and nine from *Kcns1*.

| **Gene** | **Sequence** |
|---|---|
| *Hsf4* | GGAAGGTATTAATGTTGGTATTTTTGGTTTTGTTTATGTGTTTCGGATGGTGTTTTT<br>TGTTTGTAGGTATTTGCGTTGCGAGGCGATGATAGTCGATGGCGTTCGGAAGATT<br>TGAGTCGATTGTTGGGAGAGGTG |
| *Kcns1* | CGCGTGTTGGGAGTTAGTAGTAGGCGCGACGATATTTCGAAGTTGAATTAAGCGA<br>TGTAGAAGTATTTTAGGCGGCGTAGTATCGGGTCGTCGCGTATTTTTTTGCGTTG<br>CGATTCGCGGTTATTGTAGTTATCGTCGTCGTCGTTTTTCGAGTTTGGTATTTCGG<br>TAGGTTG |

**Supplementary Table 5.** Intercept and coefficient values in epigenetic clocks 1 and 2. Coefficient value is NA for those CpG sites that have not been included in building the clock.

| | **Epigenetic clock 1** | **Epigenetic clock 2** |
|---|---|---|
| *Coefficient name* | *Coefficient value* | |
| (Intercept) | -42.60632785 | -29.3392374 |
| Gene Gm7325, position 26 | 0 | NA |
| Gene Gm7325, position 29 | 0 | NA |
| Gene Gm7325, position 38 | 0 | NA |
| Gene Gm7325, position 41 | 0 | NA |
| Gene Gm7325, position 65 | 0 | NA |
| Gene Gm7325, position 74 | 0 | NA |
| Gene Gm7325, position 95 | 0 | NA |
| Gene Gm7325, position 106 | 0 | NA |
| Gene Gm7325, position 110 | 5.97489724 | NA |
| Gene Gm7325, position 121 | 0 | NA |
| Gene Gm7325, position 135 | 0 | NA |
| Gene Gm9312, position 10 | 0 | NA |
| Gene Gm9312, position 16 | 0 | 0 |
| Gene Gm9312, position 33 | -133.0205022 | NA |

| | | |
|---|---|---|
| Gene Gm9312, position 44 | 0 | 0 |
| Gene Gm9312, position 46 | 0 | 0 |
| Gene Gm9312, position 70 | -3.05691694 | 0 |
| Gene Gm9312, position 75 | 0 | 0 |
| Gene Gm9312, position 77 | 0 | 0 |
| Gene Gm9312, position 84 | 0 | 0 |
| Gene Gm9312, position 86 | 100.1203482 | 0 |
| Gene Gm9312, position 100 | 0 | 0 |
| Gene Gm9312, position 104 | 0 | 0 |
| Gene Gm9312, position 109 | 33.27703117 | 0 |
| Gene Gm9312, position 116 | 0 | NA |
| Gene Gm9312, position 122 | 0 | 0 |
| Gene Gm9312, position 134 | 0 | 0 |
| Gene Gm9312, position 141 | 0 | 0 |
| Gene Gm9312, position 144 | 0 | 0 |
| Gene Gm9312, position 146 | 0 | 0 |
| Gene Gm9312, position 154 | 0 | 0 |
| Gene Gm9312, position 165 | 35.58595004 | 0 |
| Gene Gm9312, position 169 | 0 | 0 |
| Gene Gm9312, position 171 | 0 | 0 |
| Gene Gm9312, position 189 | 0 | 0 |
| Gene Gm9312, position 195 | 0 | 0 |
| Gene Gm9312, position 218 | -12.32714379 | NA |
| Gene Gm9312, position 224 | 0 | NA |
| Gene Hsf4, position 44 | 137.9405582 | 236.5742945 |
| Gene Hsf4, position 74 | 0 | 0 |
| Gene Hsf4, position 79 | 0 | 0 |
| Gene Hsf4, position 84 | 14.92865155 | 0 |
| Gene Hsf4, position 94 | 80.73443993 | 76.39994648 |
| Gene Hsf4, position 100 | 0 | 0 |
| Gene Hsf4, position 104 | 0 | 0 |
| Gene Hsf4, position 118 | 0 | 0 |
| Gene Kcns1, position 1 | 0 | 0 |
| Gene Kcns1, position 3 | 70.04469886 | 0 |
| Gene Kcns1, position 25 | 16.25880223 | 0 |
| Gene Kcns1, position 27 | 27.64153374 | 0 |
| Gene Kcns1, position 30 | 0 | 64.90753741 |
| Gene Kcns1, position 38 | 17.91799872 | 0 |
| Gene Kcns1, position 53 | 186.9691444 | 91.55724857 |
| Gene Kcns1, position 73 | 84.38800401 | 59.74742291 |
| Gene Kcns1, position 76 | 0 | 6.916043342 |
| Gene Kcns1, position 84 | 0 | 0 |

| | | |
|---|---|---|
| Gene Kcns1, position 89 | 0 | 0 |
| Gene Kcns1, position 92 | 0 | 0 |
| Gene Kcns1, position 94 | 0 | 0 |
| Gene Kcns1, position 107 | 0 | 88.35869557 |
| Gene Kcns1, position 112 | 0 | 0.01765114 |
| Gene Kcns1, position 117 | 205.5753337 | 0 |
| Gene Kcns1, position 119 | 0 | 0 |
| Gene Kcns1, position 135 | 0 | 0 |
| Gene Kcns1, position 138 | 29.46523008 | 0.922120902 |
| Gene Kcns1, position 141 | 0 | 0 |
| Gene Kcns1, position 144 | 36.60884345 | 2.508932843 |
| Gene Kcns1, position 151 | -136.8132736 | 0 |
| Gene Kcns1, position 165 | 239.6775681 | 57.06525829 |
| Gene Prima1, position 48 | -46.23556328 | NA |
| Gene Prima1, position 62 | 0 | NA |
| Gene Prima1, position 65 | 0 | NA |
| Gene Prima1, position 114 | 0 | NA |

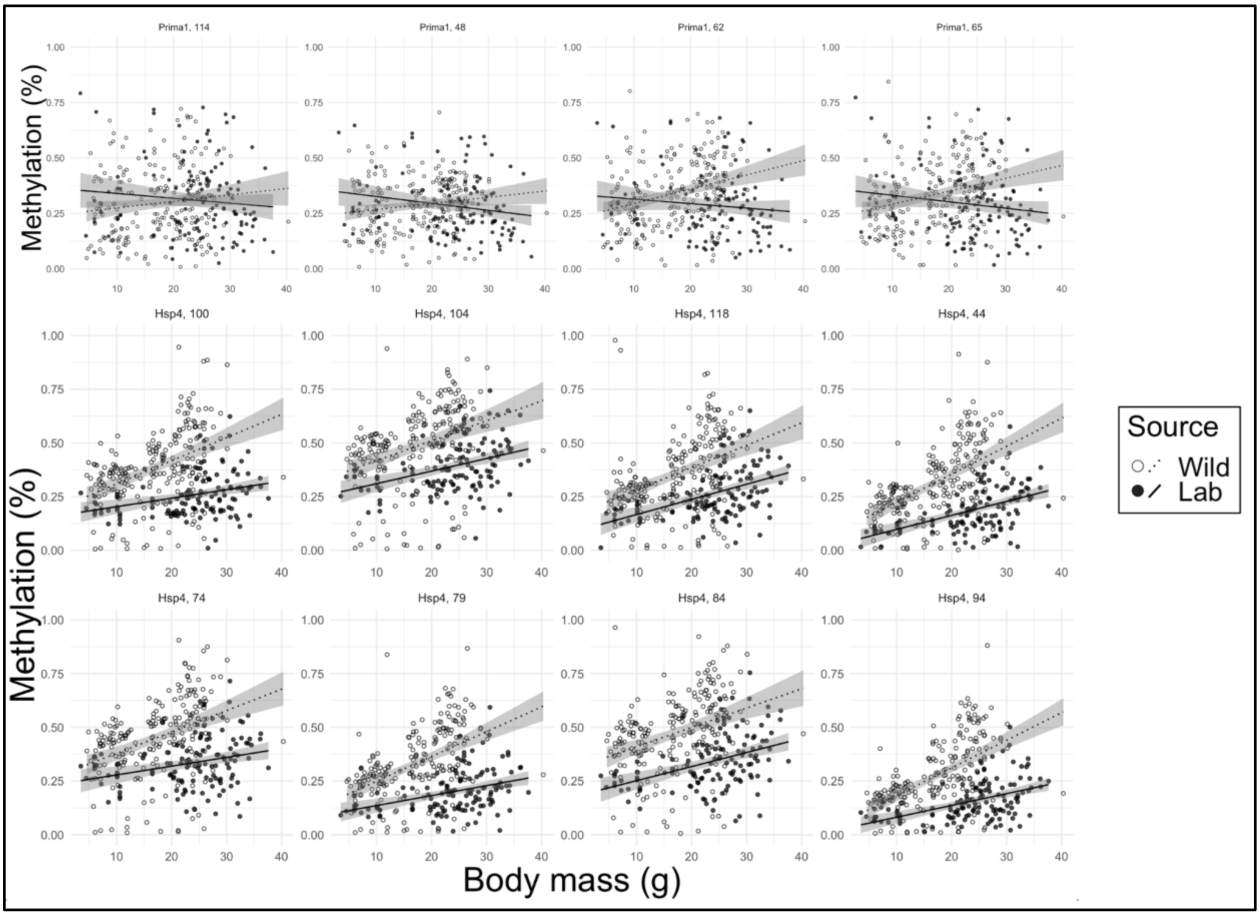


**Supplementary Figure 1. Methylation rate at 12 CpG sites from genes *Prima1* and *Hsp4* across body mass in laboratory and wild mice.** Gene and CpG start position are indicated as panel titles.

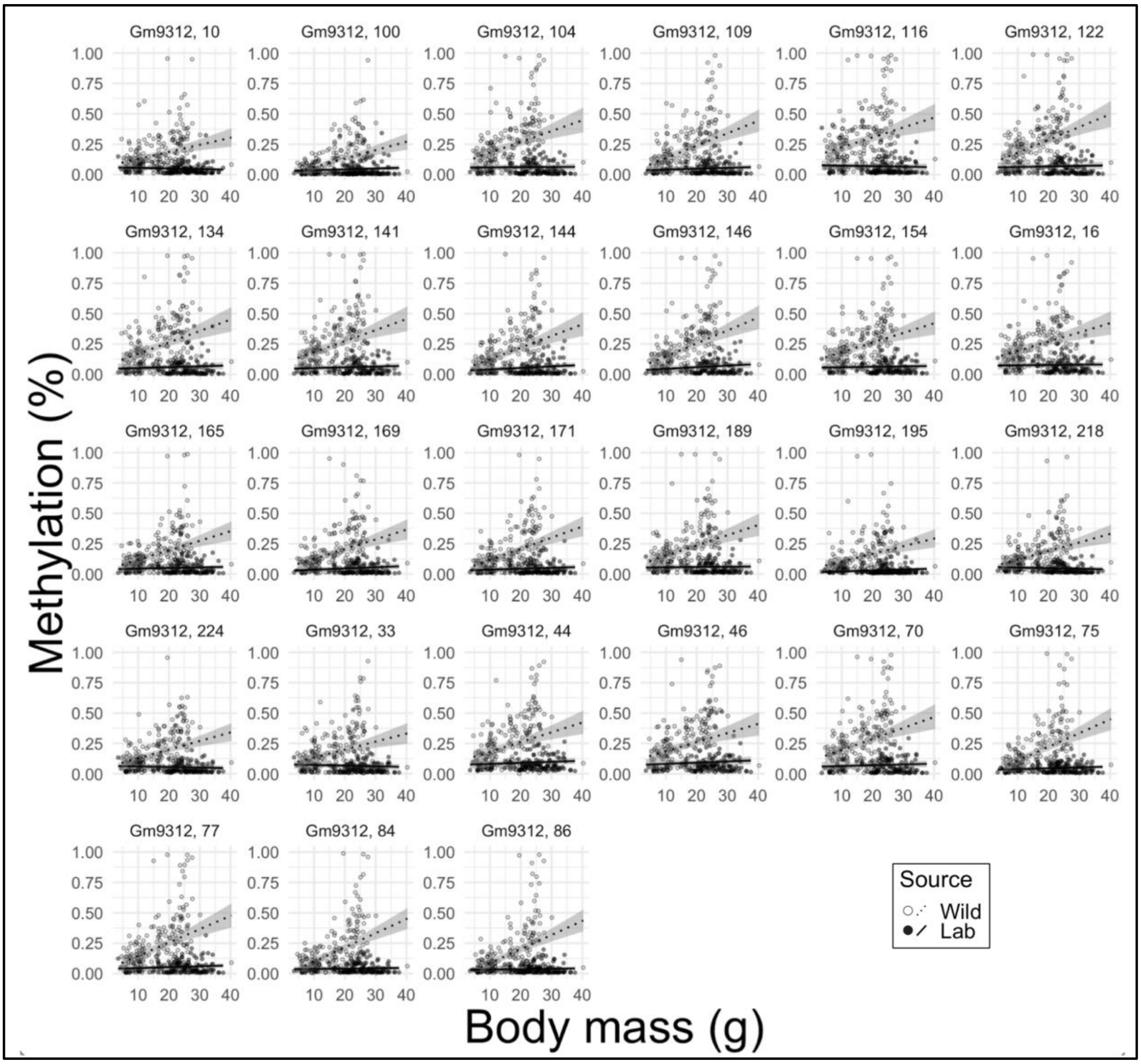


**Supplementary Figure 2. Methylation rate at 27 CpG sites from gene *Gm9312* across body mass in laboratory and wild mice.** Gene and CpG start position are indicated as panel titles.

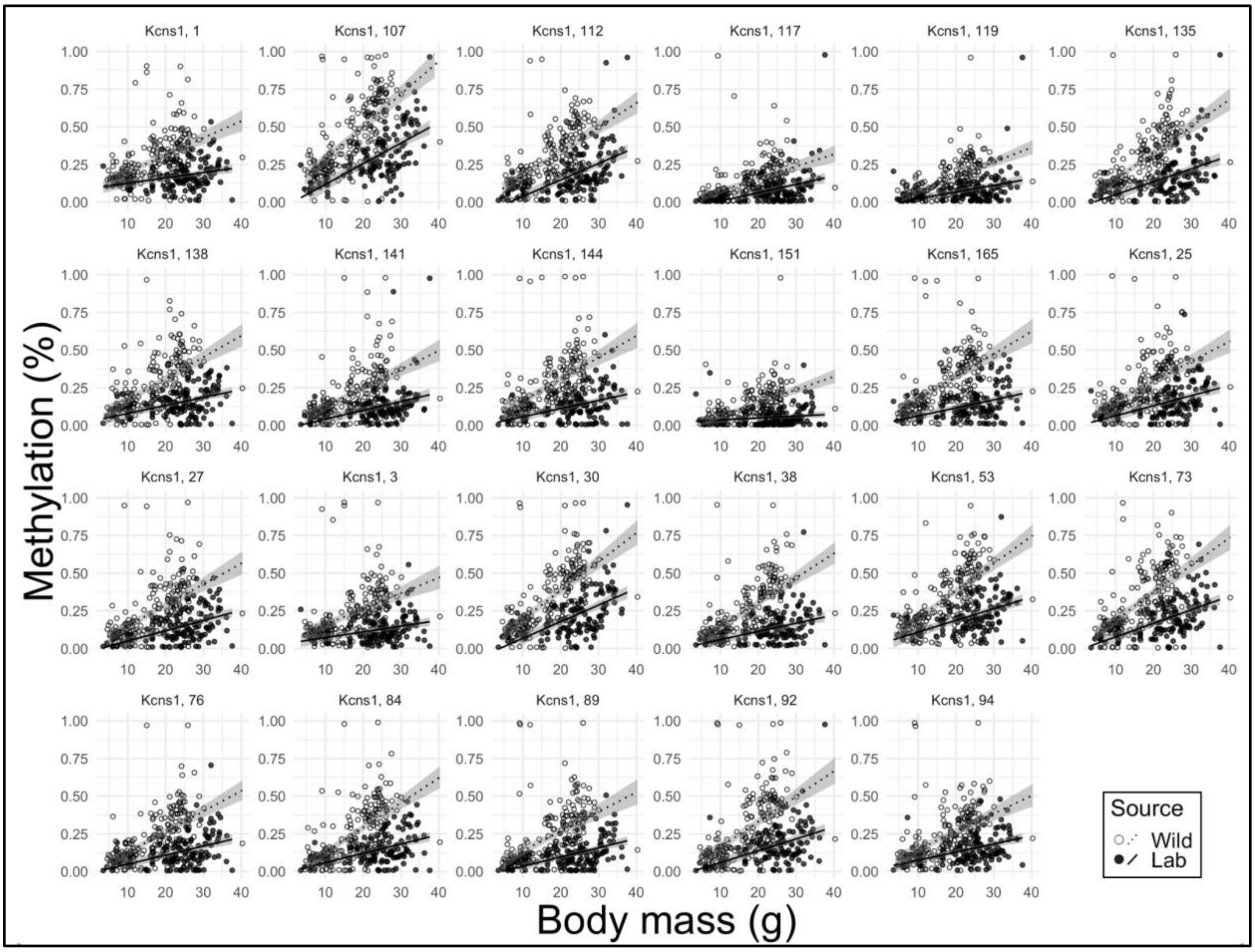


**Supplementary Figure 3. Methylation rate at 23 CpG sites from gene *Kcns1* across body mass in laboratory and wild mice.** Gene and CpG start position are indicated as panel titles.

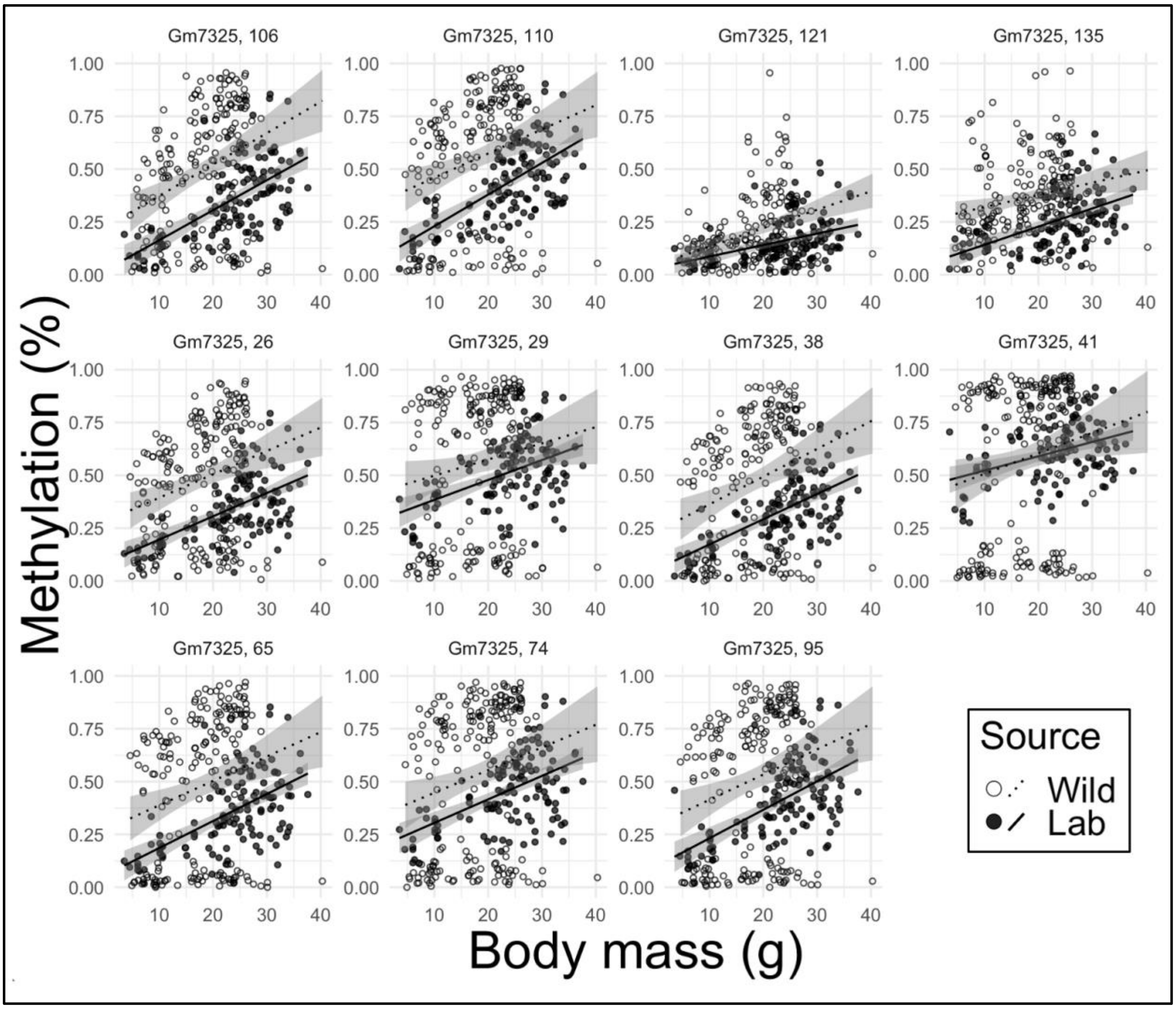


**Supplementary Figure 4. Methylation rate at 11 CpG sites from gene *Gm7325* across body mass in laboratory and wild mice.** Gene and CpG start position are indicated as panel titles.

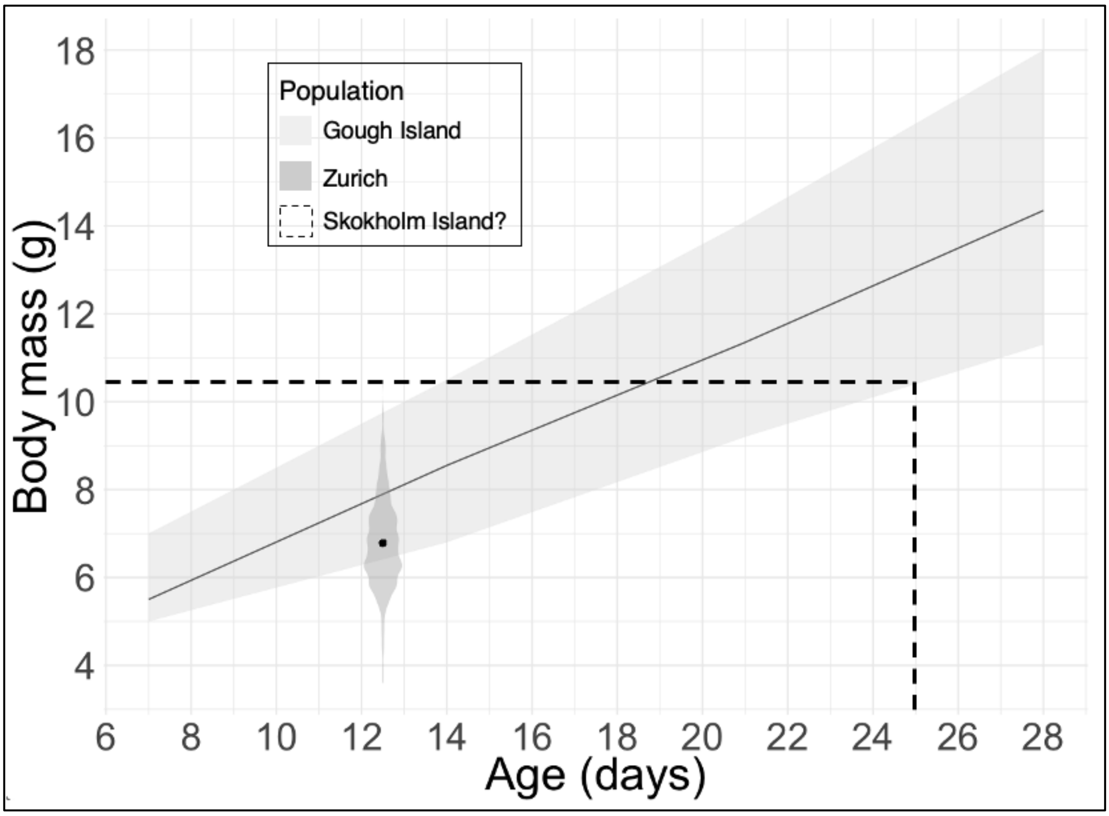


**Supplementary Figure 5. Relationship between body mass and chronological age during early life in two wild house mouse populations.** The black solid line shows the mean body mass for wild-derived (Gough Island) mice born in laboratory of a given age with shading indicating lower and upper limits (data reproduced from Figure 3 in Gray et al., 2015). The violin plot indicates body mass distribution in 12–13-day old mice (mean 12.8 days, median 13.0 days, *n*=438) from Zurich, Switzerland (Gerber et al., 2021). The point indicates average body mass in Zurich mice (6.8g; ranges 3.6–10.5g, median 6.7g). Dashed lines indicate the upper limit of body mass (10.5g) for a set of juvenile Skokholm Island mice for which epigenetic age was estimated. According to age-mass relationships in Zurich and Gough, these Skokholm Island mice are estimated to be under 25 days old.

#### References

Gerber N, Auclair Y, König B, Lindholm AK. Population density and temperature influence the return on maternal investment in wild house mice. *Front Ecol Evol.* 2021;8:602359. https://doi.org/10.3389/fevo.2020.602359

Gray MM, Parmenter MD, Hogan CA, et al. Genetics of Rapid and Extreme Size Evolution in Island Mice. *Genetics*. 2015;201(1):213-228. doi:10.1534/genetics.115.177790

Han Y, Eipel M, Franzen J, et al. Epigenetic age-predictor for mice based on three CpG sites. *Elife*. 2018;7:e37462. Published 2018 Aug 24. doi:10.7554/eLife.37462
